# Parameter and model recovery of reinforcement learning models for restless bandit problems

**DOI:** 10.1101/2021.10.27.466089

**Authors:** Ludwig Danwitz, David Mathar, Elke Smith, Deniz Tuzsus, Jan Peters

**Affiliations:** Department of Psychology, Biological Psychology, University of Cologne, Cologne, Germany

## Abstract

Multi-armed restless bandit tasks are regularly applied in psychology and cognitive neuroscience to assess exploration and exploitation behavior in structured environments. These models are also readily applied to examine effects of (virtual) brain lesions on performance, and to infer neurocomputational mechanisms using neuroimaging or pharmacological approaches. However, to infer individual, psychologically meaningful parameters from such data, computational cognitive modeling is typically applied. Recent studies indicate that softmax (SM) decision rule models that include a representation of environmental dynamics (e.g. the Kalman Filter) and additional parameters for modeling exploration and perseveration (Kalman SMEP) fit human bandit task data better than competing models. Parameter and model recovery are two central requirements for computational models: parameter recovery refers to the ability to recover true data-generating parameters; model recovery refers to the ability to correctly identify the true data generating model using model comparison techniques. Here we comprehensively examined parameter and model recovery of the Kalman SMEP model as well as nested model versions, i.e. models without the additional parameters, using simulation and Bayesian inference. Parameter recovery improved with increasing trial numbers, from around .8 for 100 trials to around .93 for 300 trials. Model recovery analyses likewise confirmed acceptable recovery of the Kalman SMEP model. Model recovery was lower for nested Kalman filter models as well as delta rule models with fixed learning rates.

Exploratory analyses examined associations of model parameters with model-free performance metrics. Random exploration, captured by the inverse softmax temperature, was associated with lower accuracy and more switches. For the exploration bonus parameter modeling directed exploration, we confirmed an inverse-U-shaped association with accuracy, such that both an excess and a lack of directed exploration reduced accuracy. Taken together, these analyses underline that the Kalman SMEP model fulfills basic requirements of a cognitive model.

## Introduction

The exploration-exploitation trade-off is the decision between a familiar option with a known reward value and an unfamiliar option with an unknown or uncertain reward value (Addicott et al., 2017). Humans face the exploration-exploitation trade-off in the context of consumer and career decisions, in decisions about how and with whom to spend their social lives, voting decisions, just to name a few examples. An individuals’ decision strategy might constitute a combination of explorative and exploitative choices or choices which vary along a continuum between exploration and exploitation (Addicott et al., 2017; Mehlhorn et al., 2015). Striking some balance between exploration and exploitation is often considered to be beneficial. Predominantly exploitative decision strategies might be disadvantageous since they might lead to inflexible, rigid and/or habitual behavior, in particular in volatile environments. An excess of exploration, on the other hand, results in inefficient switching at the expense of reward accumulation, and a lack of expertise (Addicott et al., 2017). At least two main exploration strategies have been discussed: random and directed exploration (Wilson et al., 2021). While random exploration entails randomly choosing unknown options, in directed exploration, the choices are (strategically) biased towards options that maximize information gain (reduce uncertainty). Random exploration has low computational costs but does not lead to optimal performance. Directed exploration is a more elaborate strategy involving more computational costs and leading to better performance (Meder et al., 2021; Wilson et al., 2014). Another phenomenon occurring in explore-or-exploit situations is perseveration, i.e. reward-independent choice repetition (Chakroun, 2019; Payzan-Lenestour & Bossaerts, 2012). Gershman (2020) proposes that perseveration occurs due to the trade-off between maximizing the reward and minimizing the complexity of the choice process.

To quantify behavior in the exploration-exploitation trade-off, structured environments must be created. These environments need to be structured with respect to the agent’s goal, the space of options, and the temporal dimension to ensure correct interpretations of results (Mehlhorn et al., 2015). There are several tasks providing a structured environment and enabling quantitative research on the exploration-exploitation trade-off. Among these tasks are the observe or bet task (Blanchard & Gershman, 2018; Tversky & Edwards, 1966), the patch foraging task (Constantino & Daw, 2015), the clock task (Badre et al., 2012), and multiple variants of the bandit task (Daw et al., 2006), a widely used testing bed for reinforcement learning algorithms modeled after simple slot machines with multiple arms yielding probabilistic rewards (Sutton & Barto, 1998). Bandit tasks offer a realistic and widely applicable operationalization of the exploration-exploitation trade-off, for situations in which the properties of the environment are independent of the agent’s behavior. There are various specialized versions of the bandit task, including the horizon task (Wilson et al., 2014), the leapfrog task (Blanco et al., 2013; Knox et al., 2011), and a version of the bandit task designed for testing on children (Meder et al., 2021). One task version widely used in cognitive neuroscience is the four-armed restless bandit task (Daw et al., 2006). Here, on each trial, participants select between four different colored options (“bandits”). Following selection, the number of points earned is drawn from a Gaussian distribution centered at the chosen bandit’s mean. Mean payouts of each bandit are determined by independent decaying Gaussian random walk processes.

Various models have been proposed for reinforcement learning (RL) problems such as multi-armed bandit tasks. These models generally consist of a learning rule and decision rule. The Delta rule is a model-free learning rule based on the Rescorla Wagner model (Rescorla & Wagner, 1972). Here, the chosen bandit is updated via the reward prediction error (RPE), i.e., the difference between the expected and the obtained reward, weighted by a fixed learning rate parameter (Sutton et al., 2018). Learning rates can be the same for positive and negative RPEs or differ for positive vs. negative RPEs. In what follows, the latter case is referred to as Diff Delta rule (Cazé & van der Meer, 2013). In contrast, the Kalman Filter model, also called a Bayesian Learner, is a model-based RL algorithm. In addition to tracking each bandit’s expected mean value, Kalman Filter models track the uncertainty of the expectations, and use trial-wise uncertainty-dependent learning rates. Expectations regarding unchosen options decay asymptotically towards the mean, reflecting the tasks reward structure (Chakroun et al., 2020; Daw et al., 2006; Sutton et al., 2018). Among the decision rules discussed here are the ε-greedy choice rule, and various softmax rules. The ε-greedy rule is a heuristic strategy where the agent chooses a random action in a fixed proportion of trials and chooses greedily the highest expected value (exploitation) on all other trials. The softmax decision rule (SM) also implements a form of random sampling such that options with greater expected values are chosen with higher probability. The degree to which a decision depends on option values is formalized with the softmax temperature parameter *β* (Daw et al., 2006). While values of *β* near 0 increase random exploration (for *β* = 0, choices are random), higher values of *β* correspond to increased exploitation. Additional terms can be included in the SM rule to capture relevant choice characteristics. An additional exploration bonus parameter *φ* can be implemented to grant an exploration bonus to highly uncertain and thus informative options. Higher values of *φ* correspond to increases in directed exploration (Chakroun et al., 2020; Speekenbrink & Konstantinidis, 2015). A perseveration parameter *ρ* which modulates the value of the previously chosen option can likewise be included. In principle, different learning rules can be combined with different decision rules, even though the implementation of the decision rule parameters might differ if used with different learning rules (Speekenbrink & Konstantinidis, 2015). Equations for the Delta rule, the Diff Delta rule, and the Kalman Filter, as well as the various versions of the SM decision rule are presented in more detail in the Methods section.

Behavioral measures in the bandit task, which do not require computational modeling (“model-free” measures) include the total payout, the percentage of trials in which participants chose the best bandit (“accuracy”), and the mean rank of the chosen bandit. These measures are different indicators of performance and should increase as the balance between exploration and exploitation becomes more optimal. The percentage of trials in which the participants switch from one bandit to another, on the other hand, indexes both random and directed exploration (Chakroun et al., 2020).

Several studies used model comparison to examine reinforcement learning models in human bandit task performance: Daw et al. (2006) compared three Kalman Filter models with *ε*-greedy, SM and SME decision rules. They found no evidence that a model with exploration bonus accounts better for their data. In their study the Kalman SM model outperformed the others. In line with this, Speekenbrink and Konstantinidis (2015) found the SM decision rule to outperform the SME and *ε*-greedy decision rules. With respect to learning rules, they found mixed evidence: depending on the model comparison metric, the best model used either the Kalman Filter or the Delta rule. More recently, Chakroun et al. (2020) compared all combinations of the Kalman Filter and the Delta rule with the SM, SME, and SMEP choice rules in human bandit task data. The Kalman SMEP model accounted best for their data. We recently replicated this model comparison in a group of problem gambling participants and a group of healthy matched controls (Wiehler et al., 2021). Raja Beharelle et al. (2015) found that the Kalman SME model accounted better than the Kalman SM model for a modified three-armed bandit task, which aims at preventing perseveration behavior. In other modified bandit tasks the Kalman Filter outperformed the Delta rule (Payzan-Lenestour & Bossaerts, 2012) and modeling of uncertainty-based exploration improved the fit of these models (Cogliati Dezza et al., 2017; Wilson et al., 2014). Taken together, those attempts which try to prevent or control perseveration behavior succeed in disentangling directed and random exploration.

### Aim of the Study

Even though the Kalman SMEP model was found to account best for bandit task data (Chakroun et al., 2020; Wiehler et al., 2021) parameter and model recovery work has been somewhat neglected, both by us and by others. Wilson and Collins (2019) suggest several steps researchers should follow to ensure the reliability and interpretability of computational modeling studies: first, an experiment or task has to be designed. Second, a model must be developed. The experiment and the model need to be theoretically capable of answering the research question. If this is given, third, researchers should check for model recovery and parameter recovery. As a fourth step, researchers should start the experiment and collect data. Here we use simulations to examine parameter and model recovery of the Kalman SMEP model as well as several other candidate models for human bandit task data.

Model parameters should be identifiable. That is, if the ground truth (i.e., the parameters that have produced the data) is known, it should be possible to recover these parameters using parameter estimation. This property of a model is referred to as parameter recovery. Non-deterministic models do not recover the true underlying parameters perfectly. Therefore, one aim in computational modeling is to ensure that parameter recovery is good enough for the parameters to be meaningful (Wilson & Collins, 2019). Parameter recovery is typically performed using simulated data where the true parameter values are known. Following model estimation, the true parameter values are compared to the recovered (fitted) parameter values. There is no general standard for parameter recovery and no commonly applied cut-off value, since the desired accuracy depends on the type of model and the field of research. Parameter recovery can be graphically examined as the scatter plot of the true vs. recovered parameter values to examine the degree to which they are correlated, and whether there is a bias. Additionally, ranges of the true parameter values in which parameter recovery is better or worse can be identified. In a best-case scenario, the parameter recovery does not introduce any interdependencies between the different parameters of a model. This can be examined by plotting the posterior means and deviations of the different parameters against each other (Wilson & Collins, 2019).

We also examined how parameter recovery relates to the number of data points (trials): one consideration when administering a survey or test to participants is whether performance is affected by fatigue (VandenBos, 2015). Mental fatigue can reduce behavioral flexibility and increase perseveration (van der Linden et al., 2003). Thus, mental fatigue might constitute a confounding variable in studies applying the bandit task, and hence, there should not be more trials than necessary. However, too few trials might reduce the ability to estimate the true underlying parameter values. Examining parameter recovery for different numbers of trials can therefore guide researchers to find the optimal number of trials to use in their experiments.

In a second step, we examined how model-free performance metrics relate to the data-generating parameter values. This can be helpful when evaluating psychological interpretations of model parameters. Following the conception of the exploration-exploitation trade-off (Addicott et al., 2017), both an excess of exploration as well as an excess of exploitations should negatively impact performance metrics, i.e., decreased overall payoff, lower mean rank of the chosen bandit and lower percentage of trials in which the best bandit was chosen. Random and directed exploration (softmax temperature, exploration bonus) should lead to more switches. Random exploration and perseveration are constraints to the rational solution of the exploration-exploitation trade-off (Gershman, 2020). They are therefore expected to be associated with lower performance. The exact forms of these associations are explored here.

Model recovery is given when in a set of possible models, the model underlying the data-generating process can be successfully identified using model comparison techniques. This can be examined by simulating data based on a set of different candidate models. Each simulated data set is then again fitted using the same set of candidate models. Successful model recovery then entails that the true underlying model accounts for the data better than alternative models. Crucially, model recovery is conditional on the models compared to the model of interest such that a consideration of additional models in the model recovery analysis might yield different results. Therefore, analyses of model recovery should take a range of different models into account (Wilson & Collins, 2019). Model recovery also depends on the range of input parameters used in the simulations. It is possible that model comparison can successfully distinguish models in one range of input parameters, but not in another part of the parameter space. Omitting model recovery checks can lead to misinterpretations of model comparisons and might lead to the selection of models with poor generative and predictive performance (Palminteri et al., 2017). If model recovery fails in studies of simulated data, model comparisons of models fit to empirical data are suspect, such that inferences regarding latent processes underlying cognitive functions are not interpretable. Here we analyze model recovery of Kalman Filter and Delta rule models, using a range of different variants of the softmax choice rule.

## Methods

### Material

The simulations and the analyses were conducted using the software R, version 4.0.3 (R Core Team, 2021) and Stan, version 2.21.2 (Stan Development Team, 2021). Stan is a free and open-source program that performs Bayesian inference or optimization for arbitrary user-specified models (Gelman et al., 2015). Model comparison was conducted using the R package loo (Vehtari et al., 2020). For the presentation of results, the R packages corx (Conigrave, 2020) and papaja (Aust & Barth, 2020) were used. Materials and models as well as analysis code are accessible on the website of the Open Science Foundation: https://osf.io/2e69y/.

### Underlying Payoff Structure

As input to the simulations of behavior in the four-armed bandit task, for each of the options, a decaying random walk was implemented, based on the procedure described in Daw et al. (2006). This process is specified by equations (1) and (2):

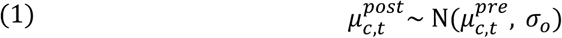

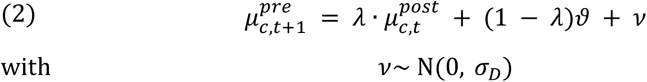

Here, the reward obtained from bandit *c* on trial 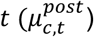 is sampled from a normal distribution with the previous reward 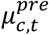 as the mean and standard deviation *σ*_*o*_. The decay of the random walk is specified in formula (2). Here, *λ* is the decay parameter, *ϑ* the center of the decay, *ν* the diffusion noise and *σ*_*D*_ the standard deviation of the diffusion noise. The following values were used as input parameters: *λ* = 0.9836, *ϑ* = 50, *σ*_*o*_ = 4 and *σ*_*D*_ = 2.8 (Daw et al., 2006). We used two different instantiations of this random walk structure (Figure 1). Analyses used the instantiation in Figure 1 A, which included 300 trials. For investigating the effect of the number of trials on parameter recovery, we used the instantiation depicted in Figure B, which included 500 trials.

**Figure 1.**
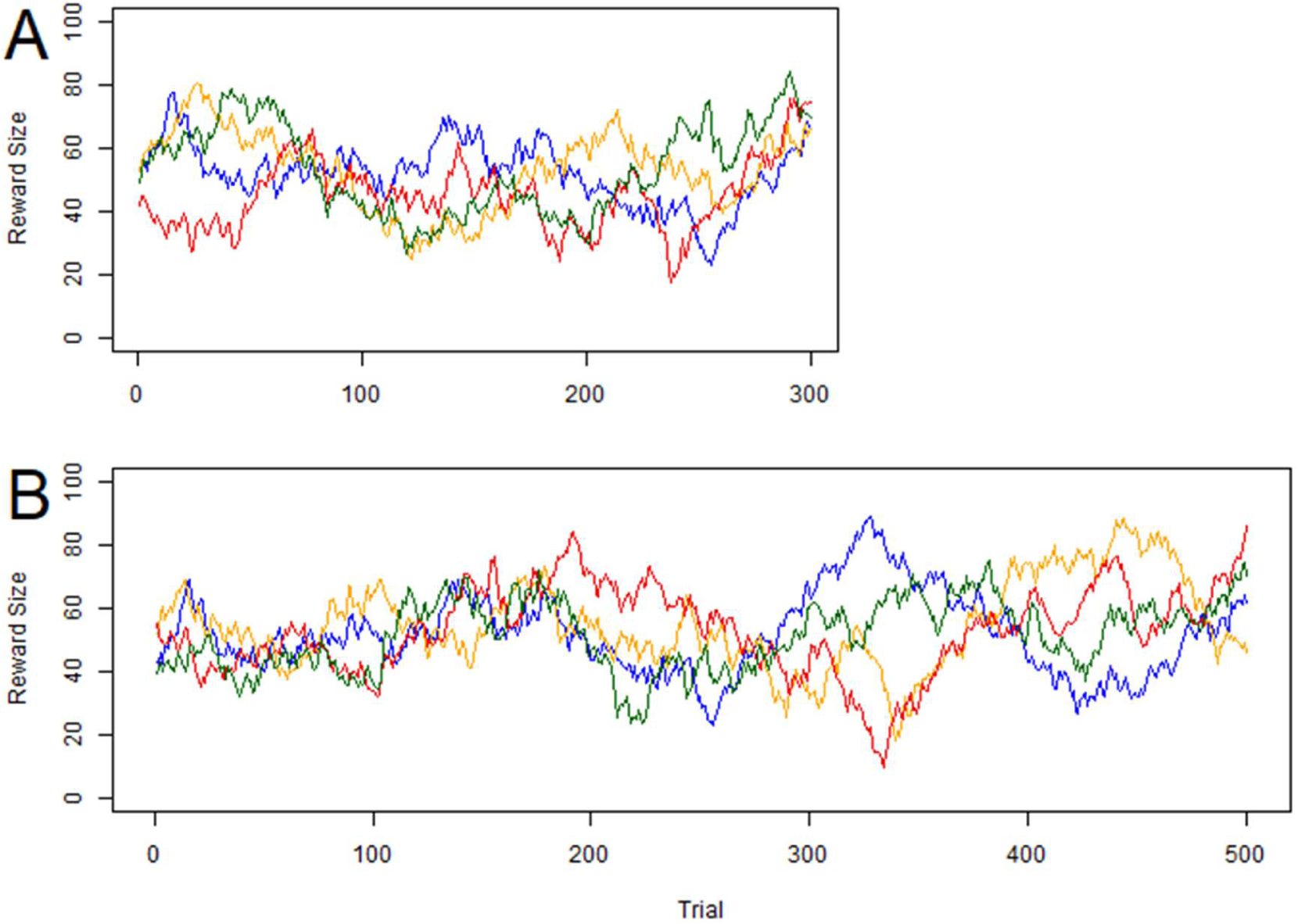
Gaussian random walks used for simulations. A: instantiation of the decaying random walk that was used in the general analysis of parameter and model recovery; B: instantiation of the random walk used to compare the parameter recovery of models with different numbers of trials. Lines reflect mean payoffs per option (bandit).

### Computational Models

To verify for model recovery, models must be compared to a set of competing candidate models (Wilson & Collins, 2019). Here, the Kalman SM, Kalman SME, Delta SM and Diff Delta SM model were compared. The Kalman SMEP model accounts for model-based RL and decision processes. It consists of a learning rule, a decision rule, and a decay rule. The learning rule specifies the updating of for the mean and variance of the expected reward based on the prediction error and an uncertainty-dependent trial-wise learning rate (Kalmain gain). It is depicted in formula (3) to (5).

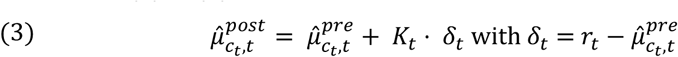

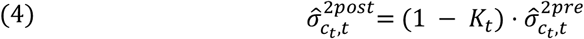

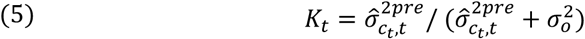

The Kalman Filter assumes that agents model a distribution for each option which assorts a credibility to each possible reward, where the mean of this distribution resembles the value of reward which appears most credible to the agent. The variance of this distribution resembles the uncertainty of this expectation. In formula (3), 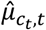 is the mean expected value of the chosen option *c*_*t*_ on a trial *t*, which is updated on each trial based on the prediction error *δ*_*t*_ and the Kalman gain *K*_*t*_. Importantly, as it is formalized in (4), not only the expected value of the chosen option is updated, but also the uncertainty of that option, i.e., the expected variance of the expected value 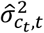. Kalman gain depends on the variance of the chosen option and the observation variance 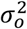, as it is shown in Equation (5). Intuitively, learning increases (*K*_*t*_ increases) with increasing option uncertainty.

The decision rule is outlined in equation (6):

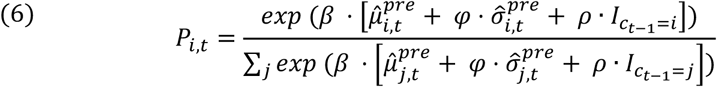

The agent chooses option *i* on trial *t* with the probability *P*_*i,t*_. This probability is calculated via softmax function with the free parameters: *β, φ* and *ρ. β* is the softmax temperature and is bounded to take only positive values (Daw et al., 2006), *φ* and *ρ* model the exploration bonus and perseveration bonus, respectively, which are implemented as additive components to the mean expected value of each option. The parameter ranges of *φ* and *ρ* are theoretically unrestricted. The exploration bonus is calculated as the product of *φ* and the standard deviation 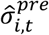 of each option *i* on trial *t*. The perseveration bonus is calculated as the product of *ρ* and an indicator function *I* that equals 1 for the option that was chosen in the previous trial and 0 for all other options. The denominator of the softmax function contains the sum over all option values.

In terms of exploration/exploitation behavior, *β* reflects the degree to which an agent uses random exploration. Small values of *β* reflect increased random exploration, whereas high values of *β* reflect a greedy strategy, in which the option with the highest expected value plus additive components is chosen deterministically. Thus, *β* reflects the trade-off between random exploration and exploitation. The range of *β* was additionally restricted on the upper bound to a maximum value of *β*_*max*_ = 3. This was done to enhance the performance of the model during the sampling process. Empirically, ranges between 0.18 and 0.26 of *β* were observed in human data (Chakroun et al., 2020). Thus, this restriction does not constrain the interpretability of the modeling results. *φ* reflects the degree of uncertainty-based exploration an agent uses: higher values of *φ* reflect higher levels of directed, uncertainty-based exploration, smaller values of *φ* reflect reduced uncertainty-based exploration. Negative values of *φ* reflect a strategy that gives an extra bonus to more certain options (Daw et al., 2006). *ρ* reflects choice “stickiness”, i.e. how much an agent tends to choose the same option as on the previous trial. High levels of *ρ* reflect a strategy strongly based on perseveration, values near zero reflect a strategy independent of the last trial and negative values reflect a value independent switching bonus (Chakroun, 2019).

The Kalman Filter SMEP model contains decay rules for the updating of the expected values and variances of all options between trials, specified in formula (7) and (8):

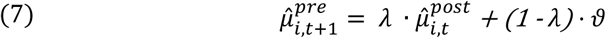

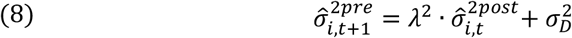

Here, the decay of the expected value depends on *λ*, which reflects the size of the steepness of the decay, while *ϑ* reflects the decay center, i.e., the value towards which the expected values decay asymptotically. Similarly, the decay of the expected variance of all options depends on *λ* and *σ*_*D*_ (Daw et al., 2006). The form of these decay functions implements that older information on the value of an option loses its validity, and the agent uses rather information about the general size of rewards of all options than rely on old information about a specific option. Uncertainty increases if there is no new information but reaches an asymptotical ceiling. In all analyses conducted on Kalman Filter models in this study, the parameters *λ, ϑ, σ*_*o*_ and *σ*_*D*_ were fixed to the true values underlying the payoff structure of the simulations. Specifically, *λ* = 0.9836, *ϑ* = 50, *σ*_*o*_ = 4 and *σ*_*D*_ = 2.8, following the implementation of Daw et al. (2006) and Chakroun et al. (2020). Similarly, the initial values *μ*_*j*,1_ and *σ*_*j*,1_, were fixed to their true values: *μ*_*j*,1_ was set to 50 and *σ*_*j*,1_ was set to 4.

The Kalman SME and the Kalman SM Model rely on the same learning and decay rules as the Kalman SMEP model (see Equations 3-8). The decision rule of the Kalman SME model resembles the Kalman SMEP model without the perseveration term. It contains the free parameters *β* and *φ*. Its decision rule is specified in formula (9).

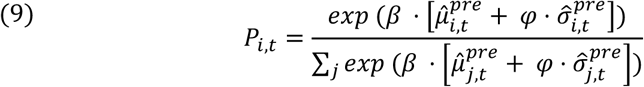

The Kalman SM Model, accordingly, resembles the Kalman SMEP model without exploration bonus and perseveration. Its only free parameter is the softmax parameter *β*. Its decision rule is specified in equation (10).

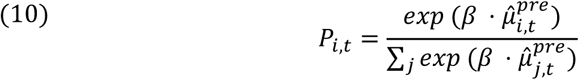

The Delta rule is a model-free learning rule, in which learning depends on the learning rate *α*. Each subject has a fixed learning rate that can take values between zero and one.

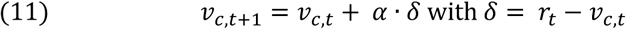

Here, *υ*_*c,t*_ is the estimated value of the chosen arm *c* on a trial *t, α* the learning rate, *δ* the prediction error and *r*_*t*_ the obtained reward. The values of unchosen options are not updated (Sutton et al., 2018). The expected values are updated with the learning rate *α*, independent of whether the prediction error is positive or negative. Initial option values *υ*_*j*,1_were set to 50 for all options. Based on the learned expectations the agent chooses an option following the softmax decision rule.

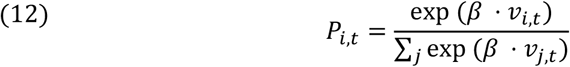

Finally, the Diff Delta rule corresponds to the Delta rule, but allows for asymmetric updating when prediction errors are positive versus when prediction errors are negative (Cazé & van der Meer, 2013). Its learning rule is described in formula (13).

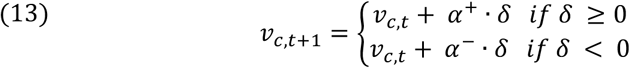

Learning rates *α*^+^ and *α*^-^ are used, depending on the sign of the prediction error. This differentiation takes into account, that humans perceive positive and negative values as distorted subjective utilities, and this distortion depends on whether the values are positive or negative (Cazé & van der Meer, 2013; Kahneman & Tversky, 1979). The decision rule of the Diff Delta SM model is the same for the Delta SM model, see equations 13. The expectations about unchosen options are, as in the Delta SM model, not updated. The Diff Delta SM model then contains three free parameters, *α*^+^, *α*^-^, and *β*, and one fixed parameter *υ*_*j*,1_, which was set to 50. An overview of the set of models and their free parameters is provided in Table S1 in the supplements.

### General Procedure of Simulation and Fitting

The process of simulation included, in each analysis, the specification of the used model, setting a range of input parameters, the specification of the underlying payoff structure and the definition of the numbers of trials and subjects to be simulated. For model estimation, the simulated data (choices, rewards) for each simulated subject and trial was entered into Stan. In Stan, the Hamiltonian Monte Carlo algorithm is used as a Markov Chain Monte Carlo method (MCMC). MCMC approximates the posterior distribution of the free parameters of the model. In contrast to frequentist approaches, where only maximum likelihood point estimates for the combination of model parameters accounting best for a certain observation are obtained, the Bayesian approach results in distributions which assort a likelihood to every value of a parameter. The width of a parameter’s posterior distribution then directly corresponds to the probability of different parameter values, given the prior and the data. The posterior distribution can also be used to estimate intervals which most likely contain the true parameters, so-called highest posterior density intervals (HDI).

We ran four chains. Chain convergence was assessed using the 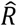 statistic, where we considered values of 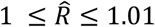 acceptable. We additionally report the effective sample size, 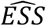, which estimates the quality of the fitting process (Kruschke, 2015). Since the first iterations in a MCMC are highly biased, a certain number of iterations in the beginning are discarded. This is called the warmup or burn in period. We used 1000 burn-in iterations and retained a further 1000 samples for analysis. No thinning was applied. For *β*, the prior was limited to the range 0 < *β* < 3; for *φ* and *ρ*, uninformed priors were used, i.e., a uniform distribution with range − ∞ to + ∞.

Model comparison was performed using the *loo* package in R, which uses a version of the loo estimate that was optimized using Pareto smoothed importance sampling (PSIS) (Vehtari et al., 2017). *loo* estimates the out-of-sample predictive accuracy of the model, i.e. how well the entire dataset without one data point predicts this excluded point.

### Procedure for Parameter Recovery

To evaluate parameter recovery, a dataset with 125 subjects and 300 trials per subject was simulated. In the following, the number of subjects is referred to as *nSubjects*, and the number of trials as *nTrials*. Wilson and Collins (2019) propose to adjust the input values of simulations to empirical obtained behavioral results. Therefore, the input values for the Kalman Filter models were adjusted to the empirical data obtained in the placebo condition by Chakroun et al. (2020). To obtain input values of *β, φ* and *ρ*, for each of the parameters, 125 values were drawn randomly from normal distributions fitted after their results. The true values of these distributions are reported in Table 1. These values resemble the values Chakroun et al. (2020) obtained in the placebo condition of their study. Here, the distribution of *β* was truncated to 0.03 < *β* < 3, hence more extreme small values indicate an entirely random choice behavior, while bigger values than 3 indicate an entirely greedy strategy. Both strategies resemble extremes of behavior in the exploration-exploitation trade-off and do not allow further examination and differentiation between subjects applying this strategy. Since *σ*_*ρ*_ in the placebo condition in Chakroun et al. (2020) was very small, likely due to a shrinkage effect due to the hierarchical estimation, *σ*_*ρ*_ was estimated based on to the standard deviation of the individual-subject posterior means in the placebo condition. The random walk depicted in Figure 1 A was included as the underlying payoff structure. The choices were simulated based on specific parameter combinations. To this end, on each trial, a choice was simulated using the softmax function with choice probabilities based on the input parameters *β*_*s*_, *φ*_*s*_, *ρ*_*s*_ and the current trial-level estimates of 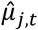 and 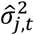.

**Table 1.** Input Values of the Simulations Used to Investigate Model Recovery

| Model | Parameters | Distribution of Input values |
| --- | --- | --- |
| Kalman SMEP | $\beta$ | $\mu = 0.23, \sigma = 0.08, lb = 0.03; ub = 3$ |
| | $\varphi$ | $\mu = 0.98, \sigma = 0.70$ |
| | $\rho$ | $\mu = 5.84, \sigma = 4.20$ |
| Kalman SME | $\beta$ | $\mu = 0.17, \sigma = 0.05, lb = 0.03; ub = 3$ |
| | $\varphi$ | $\mu = 0.15, \sigma = 0.76$ |
| Kalman SM | $\beta$ | $\mu = 0.16, \sigma = 0.06, lb = 0.03; ub = 3$ |
| Delta SM | $\beta$ | $\mu = 0.50, \sigma = 0.50, lb = 0.03; ub = 2$ |
| | $\alpha$ | $\mu = 0.50, \sigma = 0.25, lb = 0.03; ub = 1$ |
| Diff Delta SM | $\beta$ | $\mu = 0.50, \sigma = 0.50, lb = 0.03; ub = 2$ |
| | $\alpha^+$ | $\mu = 0.50, \sigma = 0.25, lb = 0.03; ub = 1$ |
| | $\alpha^-$ | $\mu = 0.50, \sigma = 0.25, lb = 0.03; ub = 1$ |

*Note.* Model Names: SM: softmax; E: exploration bonus; P: perseveration; Diff Delta means Delta rule with differential learning rates for positive and negative prediction errors. Parameters: $\beta$ is the softmax parameter; $\varphi$ is the exploration bonus parameter; $\rho$ is the perseveration parameter; $\alpha$ is the learning rate for all prediction errors; $\alpha^+$ is the learning rate that only accounts for positive prediction errors; $\alpha^-$ is the learning rate that only accounts for negative prediction errors; $\mu$ is the input true mean of the distribution of input values; $\sigma$ is the true standard deviation of the input values; $lb$ and $ub$ are the lower and upper bound of the truncated normal distribution of input values.

To test for parameter recovery, the correlations of the input values of *β*_*s*_, *φ*_*s*_ and *ρ*_*s*_ and the means of the corresponding posterior distributions were estimated. Additionally, the correlations of the input values and the size of the HDIs of the parameters were calculated to check for changes of the quality of the fitting process throughout the parameter ranges. To check for independence of the different parameters, the correlations of the obtained values of all parameters with the posterior means and HDIs of both other parameters were calculated. These associations were graphically inspected.

To inspect the influence of the number of trials on the parameter recovery, a new set of simulations was created. Due to computational feasibility constraints, in this set, *nSubjects* in each simulation 64. The generation of input values of *β*_*s*_, *φ*_*s*_ and *ρ*_*s*_ was conducted like it was depicted above. In a first step, four simulations were conducted with nTrials = 100, 200, 300 and 500. Correspondingly, the first 100, 200, 300 and 500 trials of the random walks depicted in Figure 1, B were used. These simulations were fitted using 1000 iterations in the warmup and 1000 iterations in the sampling per chain. In a second round, to examine the range between 100 and 300 parameters in greater detail, simulations with nTrials = 150, 200, 250 and 300 were conducted. Hence in the first round some 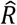 parameters were questionable, the number of iterations was set to 1500 iterations in the warmup and 1500 iterations in the sampling per chain. In a third step, the simulation with nTrials = 100 was fitted again with 1500 iterations in the warmup and 1500 iterations in the sampling per chain and prior boundaries of -10 < *φ* < 20 and -20 < *ρ* < 40.

We next calculated model-free behavioral metrics from all simulations, i.e., the percentage of switches, the total payout, the percentage of choices for the best option and the mean rank of the options. These model-free behavioral results were correlated with the input values of *β*_*s*_, *φ*_*s*_ and *ρ*_*s*_ and their associations were graphically inspected.

### Procedure of Model Recovery

The procedure of checking for model recovery included the simulation of five datasets. Each one of these was based on the Kalman SMEP model, the Kalman SME model, the Kalman SM model, the Delta SM Model, and the Diff Delta SM Model. Each of these simulations were fit by all other models. These fits were compared regarding their goodness-of-fit.

All five simulations used nSubjects = 64 and nTrials = 300. The true values of the distributions of input parameters are reported in Table 1. In line with the proposal of Wilson and Collins (2019), for the Delta Rule and Diff Delate rule modes, plausible and interpretable parameter ranges were used.

Taken together, 320 subjects, i.e., 64 subjects based on each of the five models, were simulated. Each simulated subject was fit using each of the five models. The resulting fits were compared regarding their goodness-of-fit. As a measure of goodness-of-fit, PSIS-loo was estimated (Vehtari et al., 2017). Higher loo values indicate a better fit, and the best-fitting model was determined for each simulated subject. Model recovery were then illustrated via confusion and inverse confusion matrices (Wilson & Collins, 2019). The confusion matrix quantifies the percentages of the subjects simulated based on a certain model *i* which are best fit by model *j*. Each cell contains the percentage specified in equation (14). The inverse confusion matrix quantifies the percentages of all subjects fitted best by a certain model *j* which were simulated based on a certain model *i*. Each cell contains the percentage specified in equation (15).

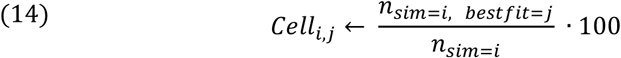

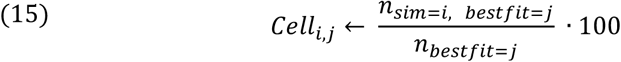

Where *sim* = *i* indicates that the subject’s data was simulated based upon model *i* and *bestfit* = *j* indicates that model *j* was found to fit the subject’s data best. The rows of the confusion matrix, which contain all subjects simulated using the same model, sum up to 100 percent, while the columns of the inverse confusion matrix, which contain all subjects fitted best by the same model, sum up to 100 percent. High values along the diagonal indicate a good model recovery while high off-diagonal values indicate poor model recovery. Diagonal values for a model in the inverse confusion matrix indicate how reliable the result of a model comparison is, if the model comparison indicated this model to fit best for a dataset (Wilson & Collins, 2019).

## Results

### Parameter recovery

To perform parameter recovery for the Kalman SMEP model, a dataset with nSubjects = 125 and nTrials = 300 was simulated and fitted. The fitting process met the requirements of representativeness, accuracy, and efficiency. The sampling parameters of all fits conducted in the analysis of parameter recovery are reported in Table S2 in the supplements.

The Pearson correlations of the true values, i.e., the input values to the simulation, and the obtained values, i.e., the means of the posterior distributions, were for *β*: *r* = .91, for *φ*: *r* = .95 and for *ρ*: *r* = .93. The scatterplots of these correlations are depicted in Figure 2: it is shown that the recovery of all parameters is generally good, while some, single values deviate from the true values. The 95% HDI is also generally rather small, indicating a concise estimation of the posterior distribution. For all parameters, the 95% HDIs include the true value. There is no apparent systematic bias. The fitting process did not introduce interdependences. For details, see Figure S1 in the supplements.

**Figure 2.**
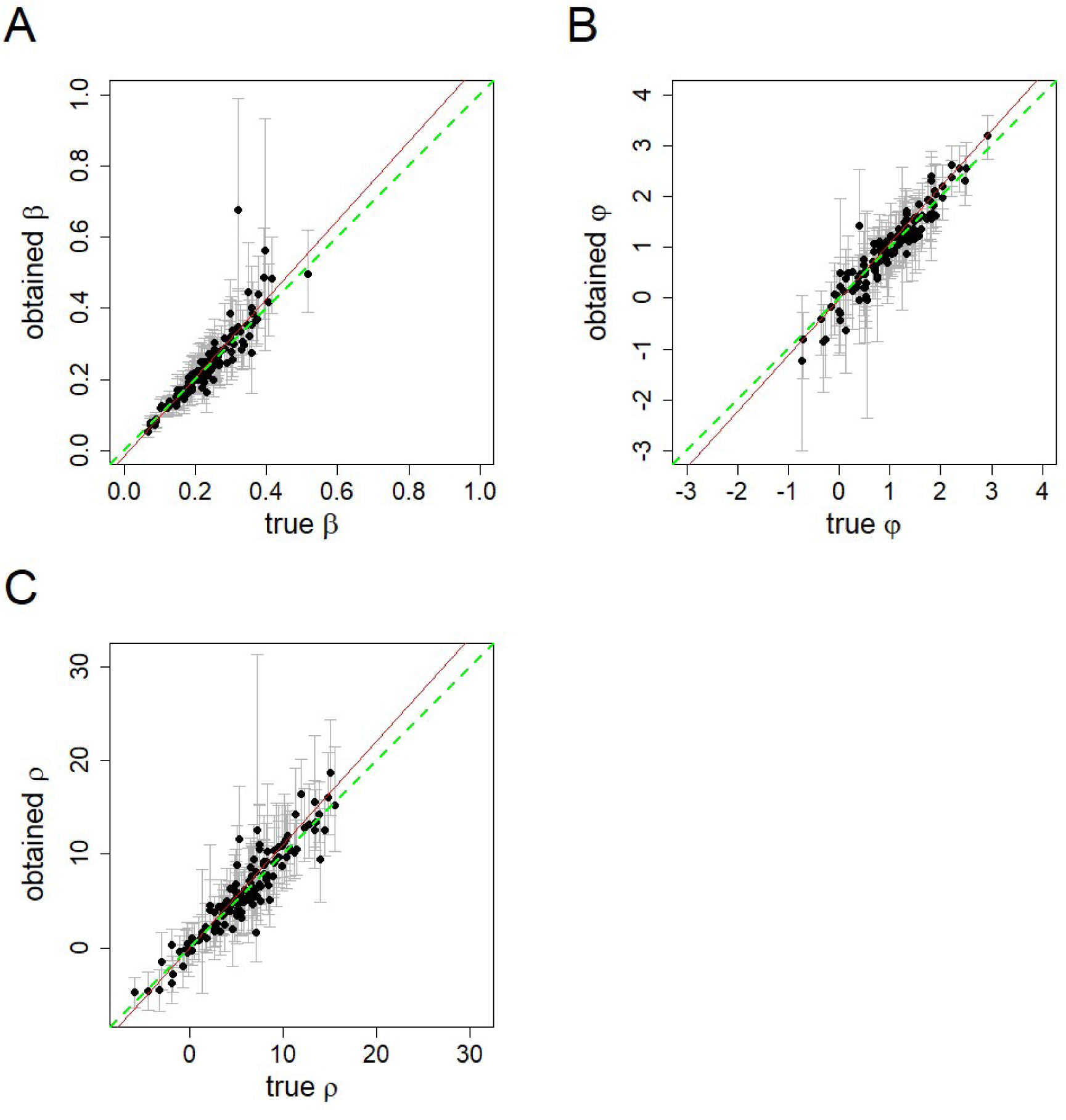
Relationship of True and Recovered Values of the Free Parameters in the Kalman SMEP Model: *β* is the softmax parameter; *φ* is the exploration bonus parameter; *ρ* is the perseveration parameter; the scatterplots show the correlation of the true values, i.e., the input values to the simulation, and the recovered values obtained in the fitting process; the mean of the posterior values are depicted as dots, the whiskers show the range of the 95% HDI; the red lines are the actual linear regression lines; the green lines show the linear regression lines for a perfect parameter recovery.

To inspect the influence of the number of trials on parameter recovery, multiple simulations and fits were created. The results of the parameter recovery for different numbers of trials are summarized in Table 2: as expected, parameter recovery improves as the number of trials increases. However, even for n=100 trials, parameter recovery was found to be acceptable.

**Table 2.** Pearson Correlations of True and Obtained Parameters, Dependent on the Number of Trials

| Parameter | Number of Trials |  |  |  |  |  |
| --- | --- | --- | --- | --- | --- | --- |
|  | 100<sup>a</sup> | 150<sup>b</sup> | 200<sup>b</sup> | 250<sup>b</sup> | 300<sup>b</sup> | 500<sup>c</sup> |
| $\beta$ | .87 | .87 | .92 | .88 | .94 | .96 |
| $\varphi$ | .82 | .87 | .89 | .94 | .92 | .94 |
| $\rho$ | .87 | .92 | .87 | .92 | .92 | .95 |

*Note.* $\beta$ is the softmax parameter; $\varphi$ is the exploration bonus parameter; $\rho$ is the perseveration parameter.
<sup>a</sup> fitting required 1500 iterations in the warmup and 1500 iterations in the sampling, the priors were limited for $\varphi$ and $\rho$ to $-10 < \varphi < 20$ and $-20 < \rho < 40$
<sup>b</sup> fitting required 1500 iterations in the warmup and 1500 iterations in the sampling
<sup>c</sup> fitting required 1000 iterations in the warmup and 1000 iterations in the sampling

**Table 3.** Bivariate Pearson Correlations of Model-Free Measures and True Values

| Measure | 1 | 2 | 3 | 4 | 5 | 6 | Mean | Standard<br>Deviation |
| --- | --- | --- | --- | --- | --- | --- | --- | --- |
| 1. True $\beta$ | - | | | | | | 0.23 | 0.08 |
| 2. True $\varphi$ | .06 | - | | | | | 1.00 | 0.69 |
| 3. True $\rho$ | -.09 | .09 | - | | | | 5.84 | 4.36 |
| 4. Payout | .65 | .12 | .17 | - |  |  | 17971.78 | 331.16 |
| 5. Percentage of Best-Option Choices | .56 | .25 | .12 | .94 | - |  | 65.38 | 6.67 |
| 6. Mean of Rank of<br>Chosen Option | .67 | .03 | .19 | .98 | .92 | - | 3.45 | 0.12 |
| 7. Percentage of<br>Switches | -.52 | .34 | -.61 | -.60 | -.42 | -.67 | 28.94 | 16.02 |

*Note.* $\beta$ is the softmax parameter; $\varphi$ is the exploration bonus parameter; $\rho$ is the perseveration parameter; true values are the input values to the simulation.

### Associations with performance metrics

Subsequently, we investigated how the free parameters of the Kalman SMEP model relate to the model-free performance metrics (Table 4, Figure 3, i.e. total payout, percentage of trials in which the best option was chosen, mean rank of the chosen bandit, percentage of switch trials). The Pearson correlations of all measures and true parameters were calculated (see Table 4). In Figure 3, the relationship of the free model parameters of the Kalman SMEP model and the model-free measures are examined more closely: higher values of *β* are associated with higher payouts and fewer switches (Figure 3A). The payout and the number of switches reach an asymptotical level, see plots A and B. Regarding *φ*, the payout obtained reaches a maximum for true values of *φ* = 1.1. Both lower and higher values of *φ* lead to a reduced payout (Figure 3C), confirming the expected inverse-U-shaped relationship between directed exploration and performance. The percentage of switches increases approximately linearly with *φ*, within the examined range of input values (Figure 3D). Regarding the effect of *ρ* on the payout, there is a slight, approximately linear growth of the payout for higher values of *ρ* within the examined range, and a strong, approximately linear decrease of the percentage of switches for higher values of *ρ* within the examined range (Figure 3E, F).

**Figure 3.**
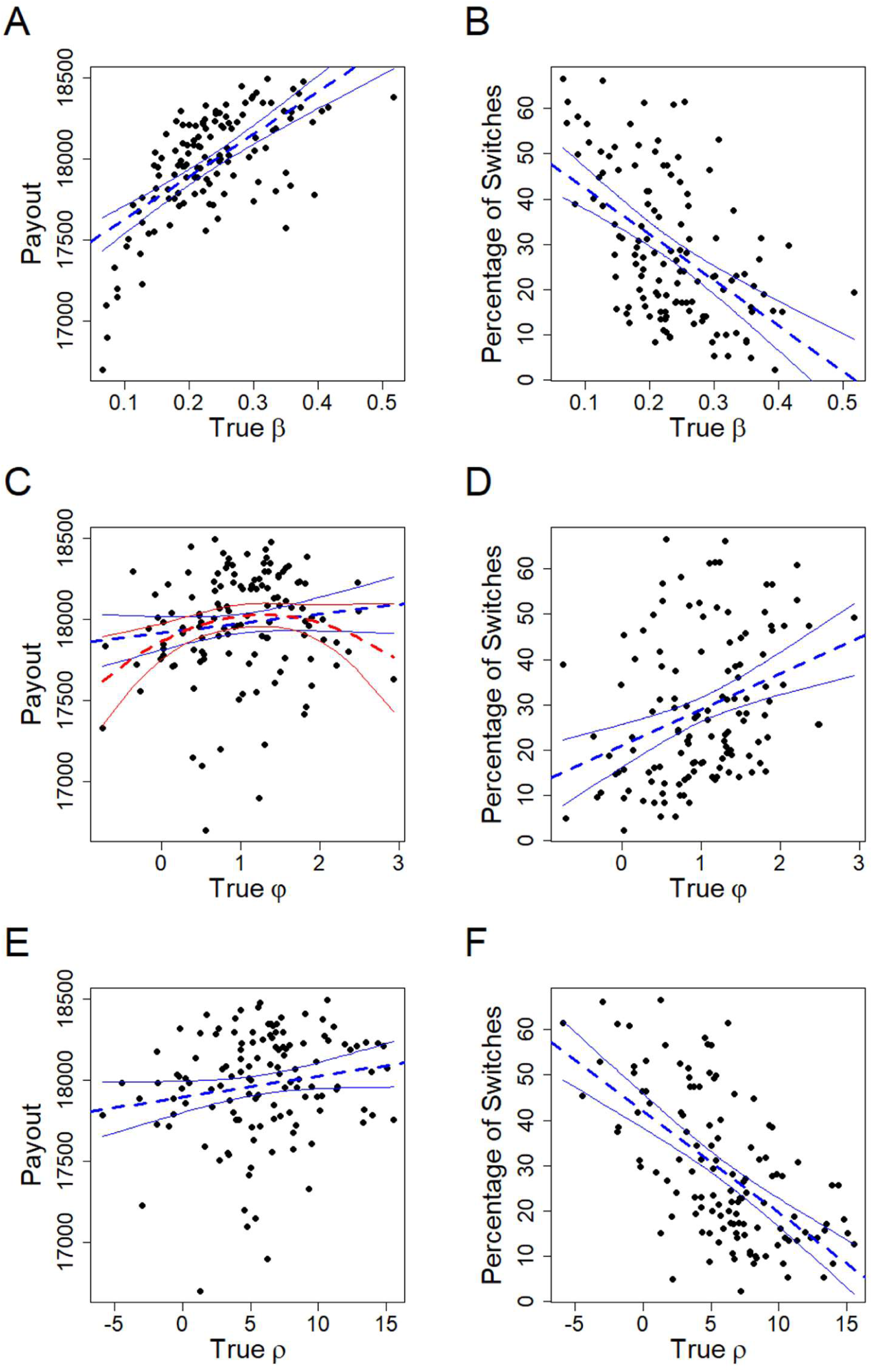
Relation of Free Parameters of the Kalman SMEP Model and Model-free Behavioral Measures: *β* is the softmax parameter; *φ* is the exploration bonus parameter; *ρ* is the perseveration parameter; true values are the input values to the simulation; payout refers to the total payout of one subject; percentage of switches refers to the percentage of trials in which another option was chosen than on the trial before; the linear regression lines and their 95% confidence interval are depicted in blue; in C, additionally the quadratic regression line and its 95% confidence interval is depicted in red.

### Model recovery

To check for model recovery, five datasets with nSubjects = 64 and nTrials = 300 each were simulated, based on the Kalman SMEP model, the Kalman SME model, the Kalman SM model, the Delta SM model, and the Diff Delta SM model. Each simulated subject was again fitted using all models and compared with respect to *loo*-based goodness-of-fit.

Figure 4 shows the confusion and inverse confusion matrices. 79.69% of subjects simulated based on the Kalman SMEP model were best fit by the Kalman SMEP model, i.e. showed successful model recovery. Some simulations were erroneously found to be fit best by the other Kalman Filter models (7.81% by Kalman SM, 10.94% by Kalman SME), whereas few were best recovered by the Delta Rule Models (1.56% by Diff Delta SM). Correspondingly, 83.61% of the simulated subjects best fit by the Kalman SMEP model were indeed simulated by the Kalman SMEP model. In some cases, the Kalman SMEP Model erroneously fitted Kalman SME simulations (13.11%) and Kalman SM simulations (3.28%). There were no cases in which the Kalman SMEP model falsely accounted for Delta Rule simulations.

**Figure 4.**
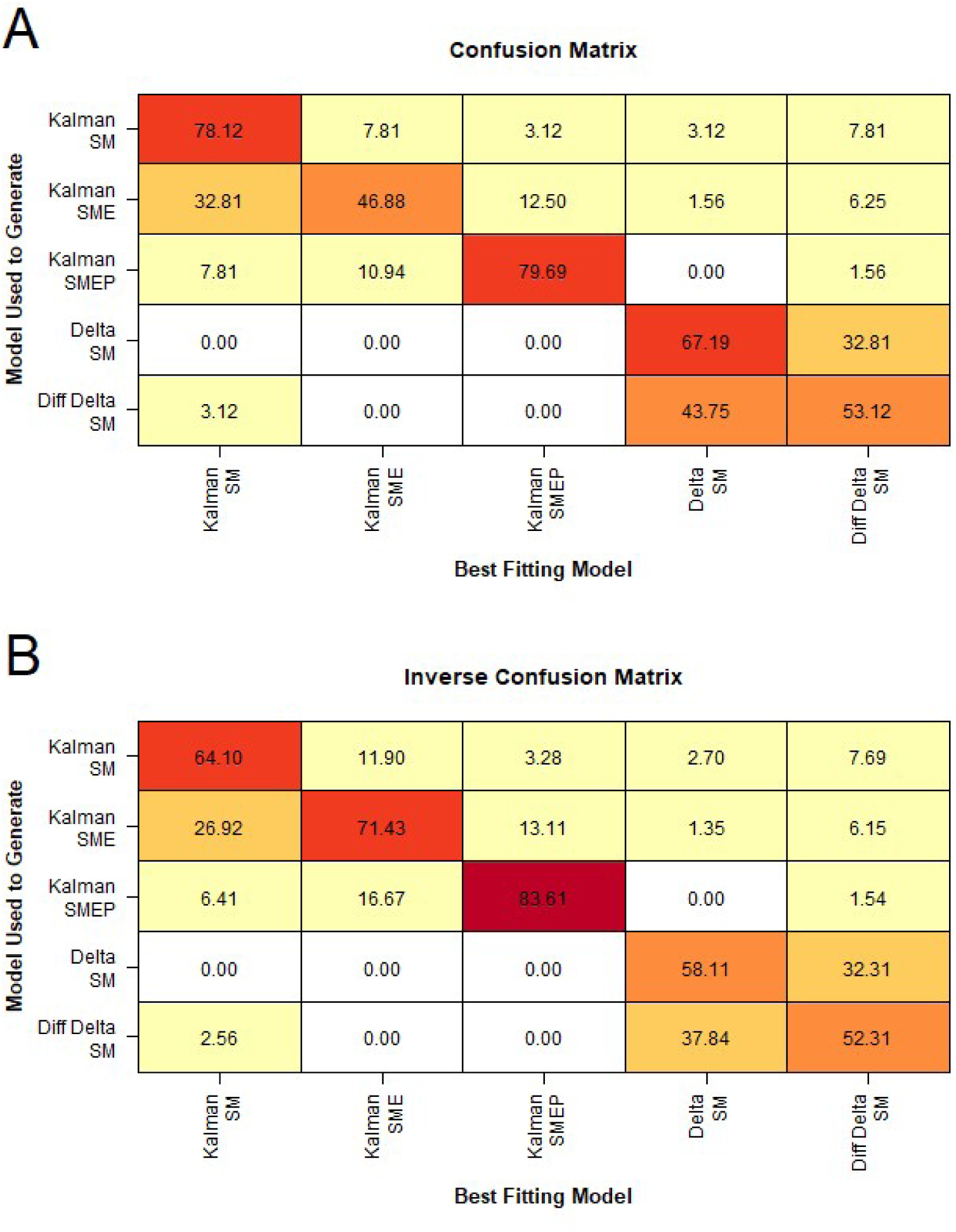
Model Recovery: Confusion Matrix and Inverse Confusion Matrix: SM: softmax; E: exploration bonus; P: perseveration, Diff Delta: Delta rule with differential learning rates for positive and negative prediction errors; the goodness-of-fit was compared using Pareto smoothed importance sampling leave-one-out cross validation; A: confusion matrix of models: percentage of all subjects simulated based on a certain model that are fitted best by a certain model; B: inverse confusion matrix: percentage of all subjects fitted best by a certain model that are simulated based on a certain model.

In addition to these overall acceptable features of the Kalman SMEP model, the confusion and the inverse confusion matrices indicate that Kalman Filter and Delta Rule models were well distinguishable from each other. In contrast, the Delta Rule SM model and the Diff Delta rule SM models are, given the range of input parameters applied, almost indistinguishable. Also, the model recovery of the Kalman SME model is questionable: only 46.88% of the simulations based on the Kalman SME model were recovered successfully and only 71.43% of the simulations the Kalman SME fitted best were really based on it. In addition to estimating the confusion matrices based on single subject’s loo values, the loo estimations of the entire datasets were compared. Based on the entire dataset, each of the simulations is fit best by the truly underlying model. See Table S3 of supplement.

## Discussion

The current study examined parameter and model recovery of reinforcement learning models for restless bandit problems. We focused on the Kalman SMEP model that has been shown to account for human data better than a range of competing (Chakroun et al., 2020; Wiehler et al., 2021), but we also examined restricted versions of that model, as well as Delta rule models, for comparison. The Kalman SMEP model combines a Kalman Filter or Bayesian Learner learning rule with a softmax decision rule with additional terms for directed exploration (exploration bonus) and perseveration. We show that the Kalman SMEP model exhibits good parameter and model recovery, even for as few as 100 trials per simulated subject. Parameters show the expected associations with model free performance metric.

For parameter recovery, correlations between true and recovered values of the Kalman SMEP model were examined, with 300 simulated trials per subject. The correlations of true and obtained values indicated a good parameter recovery (*r*′*s* >.9). The graphical inspection of the scatterplots indicated no systematic biases. Examining the influence of the number of trials of the bandit task on parameter recovery of the Kalman SMEP model, datasets were simulated with trial numbers between 100 and 500. Correlations of true and estimated parameter values generally increased with the number of trials (100 trials: .82 – .87; 500 trials: .94 – .96). Associations between true parameter values and model-free performance metrics confirmed that, for higher values of *β*, payout increased while switches decreased. For higher true values of *φ*, the frequency of switches increases, and for higher values of *ρ*, the frequency of switches decreased. We also confirmed an inverted-U-shaped association between *φ* and payout, such that both a lack and an excess of directed exploration led to performance decrements. Contrary to expectations, within the examined range of parameter values, higher values of *ρ* did not consistently result in a reduced payout.

We examined model recovery for a set of candidate models, consisting of Kalman SMEP, SME and SM models, the Delta SM model, and the Diff Delta SM model. Examination of the confusion and inverse confusion matrices showed acceptable model recovery of the Kalman SMEP model: 79.69% of simulations based on the Kalman SMEP model were recovered correctly, 83.61% of the fits accounted for best by the Kalman SMEP were in fact simulated based upon this model. Generally, model recovery performance of this model was better than for the simpler nested versions of the Kalman Filter and for the Delta rule models. The Kalman Filter and Delta rule models were well distinguishable from each other.

### Implications for Empirical Research

Our simulations have implications for empirical research applications of these models. Research using model fitting to compare parameter values between populations or conditions (Addicott et al., 2021; Chakroun et al., 2020; Wiehler et al., 2021; Zajkowski et al., 2017) can regard the correlation of true and recovered parameter values as an indicator of statistical power. Small effects might not be detected, if the accuracy of parameter recovery is not sufficient (Wilson & Collins, 2019). Also, parameter recovery sets a limit to the reliability of a measurement, such that e.g. the test-retest reliability for a given model parameter cannot exceed the correlation of true vs. recovered parameter values, even if there was a perfect temporal stability of the trait that is measured by the respective parameter.

Hence, the correlation of true and recovered variables sets a limit to the accuracy of the measurement of individual parameters, and design decisions (such as the number of trials) need to be adjusted accordingly. Even for only 100 trials, correlations of true and recovered parameter were still > .8. For studies with a large number of participants and otherwise strong manipulations or covariates this might still be sufficient. In studies with fewer participants or weaker manipulations, larger numbers of trials might be necessary to ensure adequate power.

The model-free measures of payout, the mean of the rank of the chosen option and the percentage of choices for the best option can be regarded as measures reflecting the balance of exploration and exploitation, while the number of switches only reflects overall exploration. Higher values of *β*, i.e. less random exploration, leads to a better performance (i.e. higher payout). This is in line with the conceptualization of random exploration as an inferior exploration strategy (Meder et al., 2021), which requires only little cognitive resources and is implemented in a simpler fashion into cognitive and neural processes by using neural or environmental noise to randomize choice (Zajkowski et al., 2017). Directed exploration, on the other hand, as reflected in the exploration bonus parameter *φ*, showed an inverted-U-shaped association with task performance. This resembles the theoretically-predicted inverted-U-shaped relation of exploration, exploitation, and performance, as described e.g., by Addicott et al. (2017). The disadvantages of diminished as well as excessive exploration can be observed in different psychiatric conditions: While several substance use disorders and gambling disorder are associated with diminished exploration (Morris et al., 2016; Wiehler et al., 2021), attention deficit hyperactivity syndrome and schizophrenia are associated with excessive exploration. The finding that increasing perseveration behavior, captured as *ρ*, leads to enhanced or unchanged performance in the bandit task contrasts with the conception of perseveration as a bounded rational strategy that saves cognitive resources on costs of performance accuracy (Gershman, 2020). Research mainly addressing perseveration behavior should consider this carefully. The relationship of the input parameters and the number of switches met the expectations and underlines the validity of the parameter’s implementation in the Kalman SMEP model: the percentage of switches decreases for higher values of *β* and *ρ* and increases for higher values of *φ*. This means, both exploration strategies lead to more switches, while perseveration leads to fewer switches.

The analysis of model recovery revealed that the Kalman SMEP model was distinguishable from the chosen set of candidate models. Model recovery of the Kalman SM, Kalman SME, Delta SM and Diff Delta SM model were also examined. The Delta SM and the Diff Delta SM model could be distinguished from the Kalman Filter models, whereas they were hardly distinguishable from each other. This could be a model feature, but it could also be due to the chosen range of input parameters. Still, empirical studies using model comparison to distinguish these models should be careful about the interpretation of their result.

Daw et al. (2006) used model comparison to distinguish the Kalman SME, Kalman SM and Kalman *ε*-greedy model. The Kalman SM model accounted best for the behavior of the sample (n = 14). Regarding model recovery of the Kalman SME model, even if the Kalman SME model would have been the model accounting best for the underlying decision process, the chances of Daw et al. (2006) to find this would have been limited. Chakroun et al. (2020) compared the estimates of *φ* for their empirical data between the Kalman SMEP and the Kalman SME model and found that the estimates are significantly higher in the Kalman SMEP model. This is likely due to the fact that, without a perseveration term, perseveration in the SME model is accounted for by fitting an “uncertainty-avoiding” exploration bonus parameter (Chakroun et al., 2020). Thus, our model recovery results suggest that studies might benefit from preferentially using the Kalman SMEP model rather than the Kalman SME model, even in cases in which they do not explicitly investigate perseveration behavior.

## Limitations

The current study has a number of limitations that need to be acknowledged. First, we cannot draw conclusions regarding payoff structures with different volatilities than those examined here. Similarly, parameter and model recovery of the Kalman SMEP model might differ for tasks with different numbers of arms. Regarding the effect of the number of trials on parameter recovery, a larger range of different trial numbers of trials might have been informative. Due to computational feasibility constraints, these additional simulations and fits were not carried out. Finally, is not possible to compare exhaustive sets of models in model recovery analyses (Wilson & Collins, 2019). Still, there are some shortcomings to the current analysis: The Kalman SMEP model was the most complex model of the Kalman Filter models examined. Thus, the ability to distinguish the Kalman SMEP model from Delta Rule models and more restricted Kalman Filter models was examined, while the ability to distinguish this model from other potentially more complex models was not further explored. Parameter and model recovery are dependent on the method used for the fitting. Thus, using different fitting methods like maximum likelihood might lead to different results than the here chosen MCMC approach. Empirical studies involving computational modeling often use hierarchical modeling to estimate individual parameters and group differences at once (Chakroun et al., 2020; Raja Beharelle et al., 2015; Wiehler et al., 2021).In this study, we focused on fitting single subject values. Adding a hierarchical level to the model might influence the outcomes of parameter and model recovery on the subject level.

## Conclusion

The present study examined the parameter and model recovery of the Kalman SMEP model for restless bandit problems. Parameter recover of the Kalman SMEP was excellent for 300 trials, and acceptable even for as few as 100 trials per simulated subject. Model parameters of the Kalman SMEP model showed associations with model-free measures of performance and behavior, in line with their typical psychological interpretations. Model recovery confirmed that the Kalman SMEP model was distinguishable from simpler nested Kalman Filter models as well as Delta Rule models. Future empirical studies that utilize computational reinforcement learning models in the context of restless bandit problems may benefit from the simulation work reported here.

## List of Abbreviations

BIC: Bayesian information criterion
Diff Delta rule: Delta rule with differential learning rates for positive and negative prediction errors
HDI: highest density interval
HMC: Hamiltonian Monte Carlo
loo: leave-one-out cross validation
LOWESS: locally weighted scatterplot smoothing
MCMC: Markov Chains Monte Carlo
nSubjects: number of subjects
nTrials: number of trials
PSIS: Pareto smoothed importance sampling
RL: reinforcement learning
RPE: reward prediction error
SM: softmax
SME: softmax, exploration bonus
SMEP: softmax, exploration bonus, perseveration bonus
SPE: state prediction error
TD: temporal discounting
TMS: transcranial magnet stimulation
UCB: upper confidence bound

## List of Symbols

*α*: learning rate
*α*^+^: learning rate for positive prediction errors
*α*^-^: learning rate for negative prediction errors
*β*: softmax temperature parameter (inverse random exploration)
*δ*: reward prediction error
*ϑ*: decay center
*Κ*: Kalman learning rate
*λ*: decay parameter
*μ*: mean of subjective reward probability distribution
*μ*_*β*_: mean of input values of the softmax parameter
*μ*_*φ*_: mean of input values of the exploration bonus parameter
*μ*_*ρ*_: mean of input values of the perseveration bonus parameter
*ρ*: perseveration bonus parameter
*σ*^2^: variance of subjective reward probability distribution
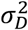: variance of decay
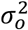: variance of observation
*σ*_*β*_: standard deviation of input values of the softmax parameter
*σ*_*φ*_: standard deviation of input values of the exploration bonus parameter
*σ*_*ρ*_: standard deviation of input values of the perseveration bonus parameter
*φ*: exploration bonus parameter
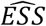: effective sample size
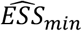: minimal effective sample size
*ϑ*: expected reward value
*r*: reward
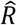: estimation for merging of MCMCs
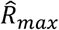: maximal 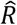, indicates most detrimental merging of MCMCs

## Supplement A: Table S1

**Table S1.** Overview of the Examined Models and their Free and Fixed Parameters

| Model | Free Parameters | Fixed Parameters |
| --- | --- | --- |
| Kalman SMEP | $\beta, \varphi, \rho$ | $\lambda, \vartheta, \sigma_o^2, \sigma_D^2, \mu_{j,1}, \sigma_{j,1}^2$ |
| Kalman SME | $\beta, \varphi$ | $\lambda, \vartheta, \sigma_o^2, \sigma_D^2, \mu_{j,1}, \sigma_{j,1}^2$ |
| Kalman SM | $\beta$ | $\lambda, \vartheta, \sigma_o^2, \sigma_D^2, \mu_{j,1}, \sigma_{j,1}^2$ |
| Delta SM | $\beta, \alpha$ | $v_{j,1}$ |
| Diff Delta SM | $\beta, \alpha^+, \alpha^-$ | $v_{j,1}$ |

*Note.* Model Names: SM: softmax; E: exploration bonus; P: perseveration, Diff Delta: Delta rule with differential learning rates for positive and negative prediction errors. Free Parameters: $\beta$ is the softmax parameter; $\varphi$ is the exploration bonus parameter; $\rho$ is the perseveration parameter; $\alpha$ is the learning rate for all prediction errors; $\alpha^+$ is the learning rate that only accounts for positive prediction errors; $\alpha^-$ is the learning rate that only accounts for negative prediction errors. Fixed Parameters: $\lambda$ is the decay parameter; $\vartheta$ is the decay center; $\sigma_o^2$ is the estimated observation variance; $\sigma_D^2$ is the estimated diffusion variance; $\mu_{j,1}$ is the initial prior mean of the expected reward for all options; $\sigma_{j,1}^2$ is the initial prior variance of the expected reward for all options; $v_{j,1}$ is the initial expected reward value for all options.

## Supplement B: Table S2

**Table S2.** Sampling Diagnostics of the Fits in Analysis of Parameter Recovery

| Simulating Model | Fitting Model | Number of Iterations | Number of Subjects | Number of Trials | $\hat{R}_{max}$ | $\widehat{ESS}_{min}$ |
| --- | --- | --- | --- | --- | --- | --- |
| Kalman SMEP | Kalman SMEP | 2000 | 150 | 300 | 1.0088 | 997 |
| Kalman SMEP | Kalman SMEP | 2000 | 64 | 100 | 5.79 | 2 |
| Kalman SMEP | Kalman SMEP<br>+ Bounds | 3000 | 64 | 100 | 1.0077 | 407 |
| Kalman SMEP | Kalman SMEP | 3000 | 64 | 150 | 1.012 | 418 |
| Kalman SMEP | Kalman SMEP | 2000 | 64 | 200 | 1.034 | 117 |
| Kalman SMEP | Kalman SMEP | 3000 | 64 | 200 | 1.030 | 136 |
| Kalman SMEP | Kalman SMEP | 3000 | 64 | 250 | 1.0037 | 1512 |
| Kalman SMEP | Kalman SMEP | 2000 | 64 | 300 | 1.0084 | 1239 |
| Kalman SMEP | Kalman SMEP | 3000 | 64 | 300 | 1.0037 | 1743 |
| Kalman SMEP | Kalman SMEP | 2000 | 64 | 500 | 1.0072 | 1197 |

*Note.* Kalman SMEP means Kalman Filter Model with Softmax decision rule, exploration bonus and perseveration bonus; Bounds marks the case in which the range of possible values of $\varphi$ was restricted to $-10 < \varphi < 20$ and the range of $\rho$ was restricted to $-20 < \rho < 40$ ; the iterations were in each case half warmup and half sampling; $\hat{R}_{max}$ the maximum $\hat{R}$ estimated for a parameter in the corresponding fit, indicating how well the chains merged; $\widehat{ESS}_{min}$ is the minimum $\widehat{ESS}$ estimated for a parameter in the corresponding fit, indicating the efficiency and accuracy of the fit.

## Supplement C: Figure S1

**Figure S1.**
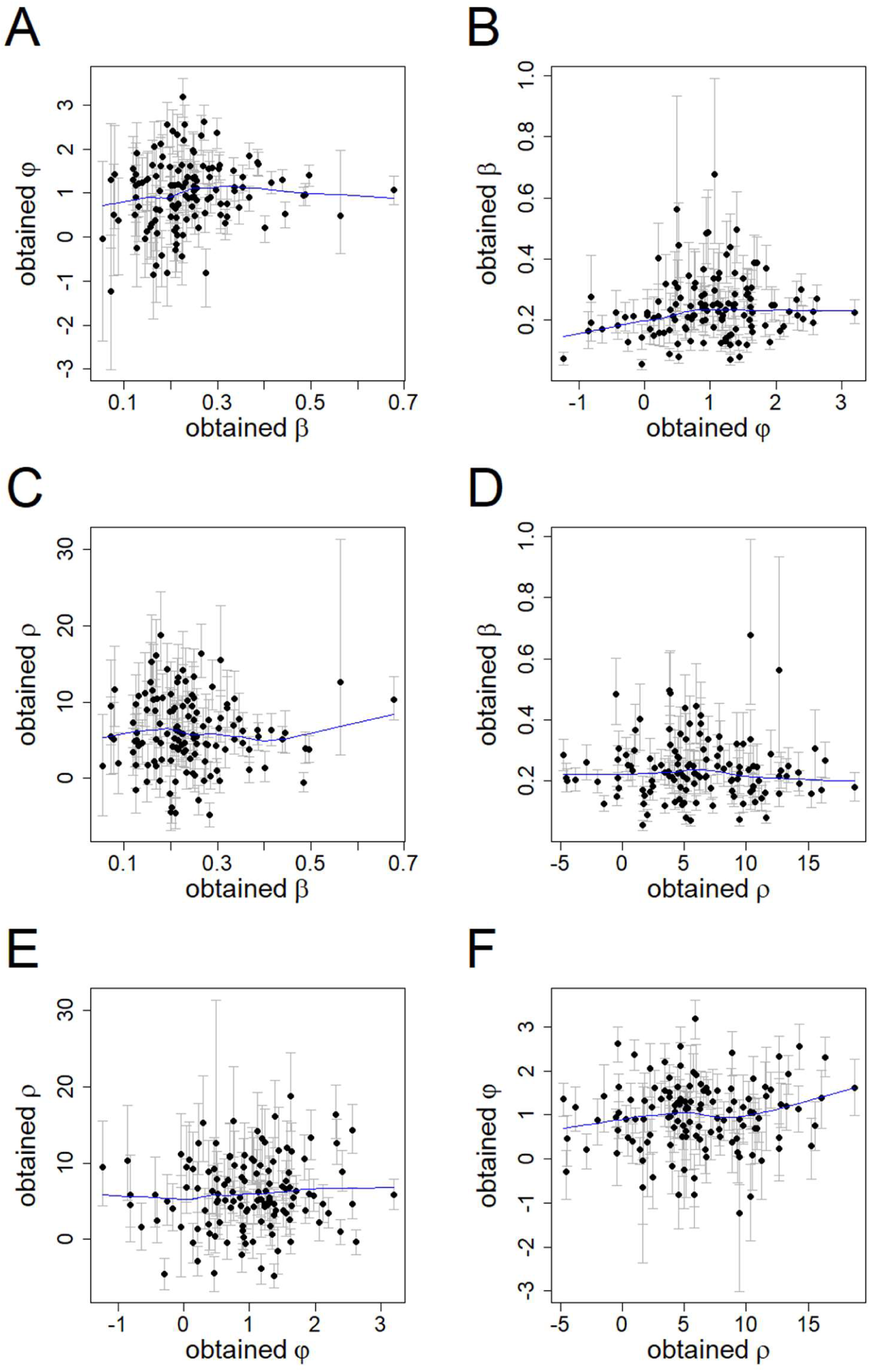
Interdependence of the Recovered Free Parameter Values of the Kalman SMEP Model: *β* is the softmax parameter; *φ* is the exploration bonus parameter; *ρ* is the perseveration parameter; the scatterplots show the correlation of the different recovered parameters; the mean of the posterior values is depicted as dot the whiskers show the range of the 95% HDI; the locally weighted scatterplot smoothing (LOWESS) lines of the relationships are depicted in blue

## Supplement D: Table S3

**Table S3.** Comparison of loo Estimates for the Entire Datasets

| Model used to fit | Model used to generate |  |  |  |  |
| --- | --- | --- | --- | --- | --- |
|  | 1 | 2 | 3 | 4 | 5 |
| 1. Kalman SMEP | -0.6081<sup>a</sup> | -0.7430 | -0.7845 | -0.2616 | -0.3402 |
| 2. Kalman SME | -0.6476 | -0.7413<sup>a</sup> | -0.7825 | -0.2596 | -0.3378 |
| 3. Kalman SM | -0.6753 | -0.7531 | -0.7801<sup>a</sup> | -0.3497 | -0.4126 |
| 4. Delta SM | -0.6866 | -0.7621 | -0.7929 | -0.2230<sup>a</sup> | -0.3109 |
| 5. Diff Delta SM | -0.6837 | -0.7617 | -0.7926 | -0.2235 | -0.3036<sup>a</sup> |

*Note.* Model Names: SM: softmax; E: exploration bonus; P: perseveration, Diff Delta: Delta rule with differential learning rates for positive and negative prediction errors. The goodness-of-fit was compared using Pareto smoothed importance sampling leave-one-out cross validation: the cells contain the estimated loo measure for the entire fit, divided through the number of subjects and number of trials.
<sup>a</sup> the best fitting model of the column.

